## Supplementary Information for "Integrated cross-study datasets of genetic dependencies in cancer"

Pacini et al.

#### Supplementary Tables

**Supplementary Table 1** - List of cell lines included in at least one of the two individual screens with cross-institute identifiers, lineage and cancer subtype annotations and dataset of origin.

**Supplementary Table 2** - Top 10 list of enriched MsigDB gene sets found for the first two principal components of the CRISPRcleanR processed dataset.

**Supplementary Table 3** - List of significant tissue specific biomarker and dependency associations under the three different pre-processing methods and four batch correction pipelines.

**Supplementary Table 4** - List of common essential genes with associated Tiers.

**Supplementary Table 5** - Top 20 list of gene ontology Biological Process gene sets from MsigDB unique to the common essential gene set of the integrated data

**Supplementary Table 6** - Lists of tissue specific biomarker/dependency associations found as significant in the integrated dataset but not the individual datasets.

### Supplementary Figures

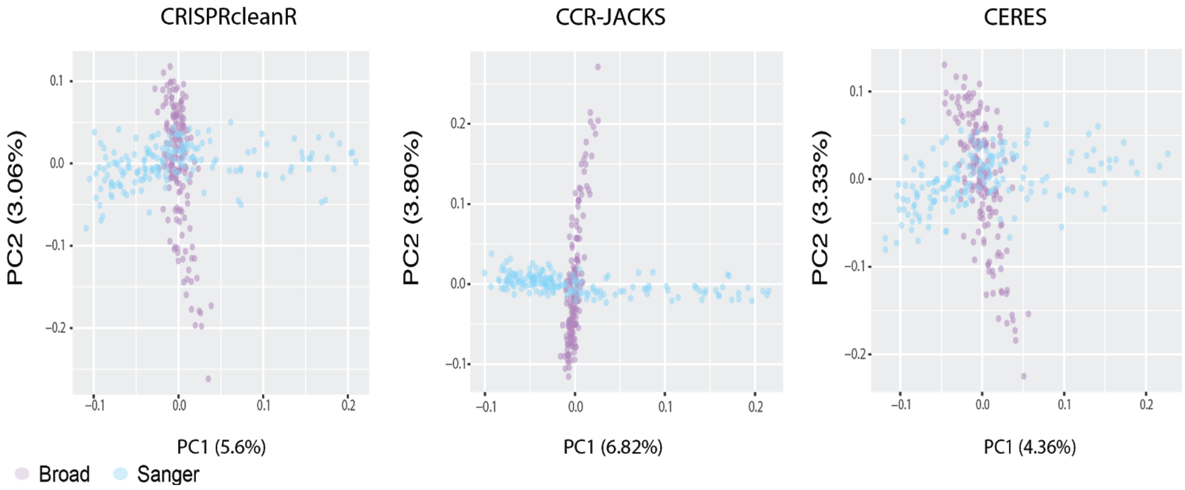


**Supplementary Figure 1: Residual batch effects following correction across pre-processing methods.** Principal component plots of the gene dependency profiles of cell lines screened in both Broad and Sanger studies (168 cell lines) following ComBat batch correction, across different pre-processing methods.

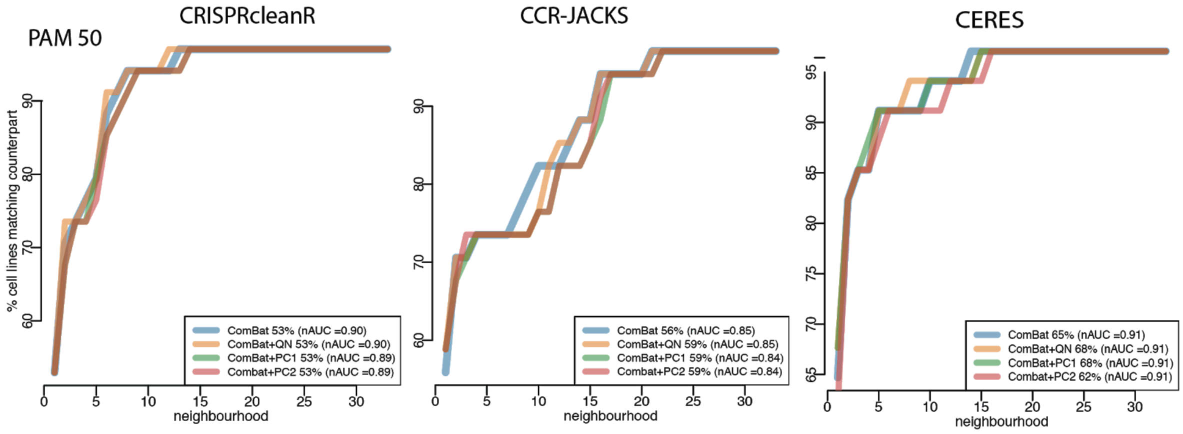


**Supplementary Figure 2: Lineage subtype identification**.

Agreement of Breast CRISPR-cas9 fitness profiles according to the clinical Breast PAM50 cancer subtypes. For each query Breast cancer cell line in turn we computed correlation scores to all other Breast cancer cell lines (responses). We then ranked the response cell lines according to these correlations. For each query cell line, the rank position k of the most correlated response cell line from the same cancer subtype (matching response) was identified. A rank of k = 1 indicates that the query cell line was closest to another cell line from the same cancer subtype. The curves show the ratio of query cell lines with a matching response with a given rank position. The proportion of query cell lines with a matching response in k = 1 are also shown as percentages for each dataset. The normalised area under the curve (nAUC) for each dataset is shown in brackets.

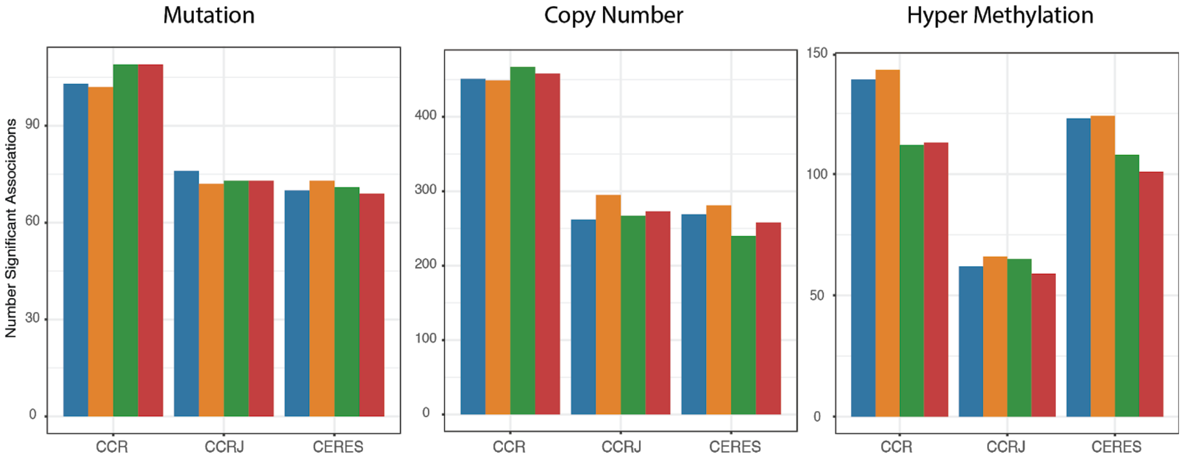


**Supplementary Figure 3: Identification of Biomarker/Dependency associations**

Cancer functional-events/dependency associations found for each biomarker type and dataset at 5% FDR.

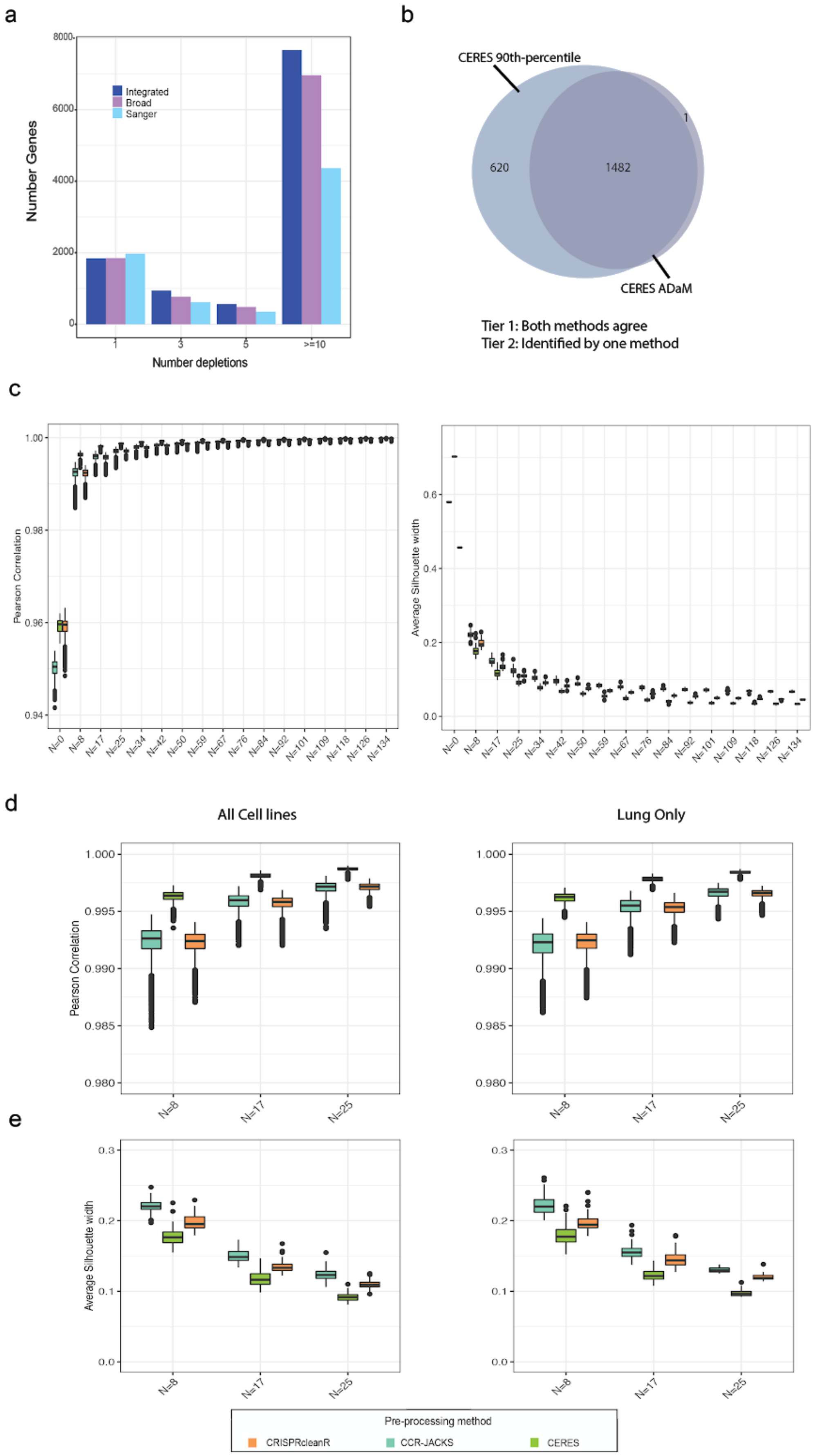

a
b
c
d
e
Number Genes
Number depletions
Integrated
Broad
Sanger
CERES 90th-percentile
CERES ADaM
620
1482
1
Tier 1: Both methods agree
Tier 2: Identified by one method
Pearson Correlation
Average Silhouette width
All Cell lines
Lung Only
Pre-processing method
CRISPRcleanR
CCR-JACKS
CERES

**Supplementary Figure 4: Performance of the Integrated Dataset.** a. Number of genes that are significant dependencies (at 5% FDR) in fixed numbers of cell lines across individual datasets and the integrated one, for CRISPRcleanR pre-processing methods. b. Venn diagram showing genes called common essential when using two different detection algorithms (ADaM and 90th-percentile) applied to the CERES processed dataset. c. The boxplots contain 50 random samples of between 5% and 90% of the 168 overlapping cell lines (number of cell lines in each sample indicated on the x-axis). For each sample the Pearson correlation of the DPGs following ComBat correction compared to the integrated dataset was calculated for each pre-processing method. We also include N=0 as the correlation between the dataset with no batch correction applied. In the right hand plot the average silhouette width (ASW) for each downsampled dataset was calculated using the institute of origin as the cluster label. An ASW of close to zero indicating a near random performance of the clustering, meaning the samples do not cluster by the origin of the screen and batch effects have been removed. d. The boxplots contain 50 random samples of 8, 17 or 25 cell lines from the overlapping set of 168 cell lines on the left plot. The right hand plot shows the results when using 8, 17 or 25 cell lines drawn only from the set of Lung cancer cell lines. For each sample the Pearson correlation of the DPGs following ComBat correction compared to the integrated dataset was calculated for each pre-processing method. e. The average silhouette width (ASW) for each downsampled dataset was calculated using the institute of origin as the cluster label. An ASW of close to zero indicating a near random performance of the clustering, meaning the samples do not cluster by the origin of the screen and batch effects have been removed. The left hand plot draws random samples of cell lines from all lineages, the right hand plot uses Lung cancer cell lines only.
